# CRISPR epi-editing reveals regulatory crosstalk between alternative promoters and splicing in Neurexin isoform expression

**DOI:** 10.1101/2025.09.10.675405

**Authors:** Yungyi Hsiao, Andrea M Gomez

## Abstract

The use of alternative promoters and splicing increases molecular complexity and diversifies cellular functions. However, mechanisms of crosstalk between transcription and splicing remain poorly understood. Here, we utilize CRISPR epi-editing in neurons to manipulate isoforms of the synaptic organizer, Neurexin-1, and elucidate mechanisms underlying the co-regulation of alternative promoters and splicing. Surprisingly, silencing individual Neurexin-1 promoters altered downstream promoter activity via transcriptional interference and biased splicing decisions. Our data reveals transcriptional interference as key to shaping cell type-specific Neurexin-1 isoforms in the mouse hippocampus and demonstrates the power of epi-editing to uncover regulatory interactions between RNA processes in the brain.

## Introduction

The high degree of alternative promoter usage and alternative RNA splicing by neurons establishes precise patterns of connectivity in the brain and contributes to phenotypic variation in vertebrates^1,2^. Transcription initiated from alternative promoters within a single gene generates diverse isoforms with varying functions ^3^. Alternative RNA splicing results from the alternate choices made during mRNA preprocessing, as introns are removed and exons spliced together. Alternative promoter and splicing choices can alter the protein function by changing ligand-binding domains, domains targeted by posttranslational modification, and intrinsically disordered domains ^4^. Notably, half of all mammalian genes have at least two promoters ^5–7^, while 95% of multi-exon genes undergo alternative splicing ^8,9^. Thus, the extensive isoform generation through alternative promoter and splicing in neurons provides the molecular complexity necessary for establishing and specializing synaptic properties ^10–12^.

Genes that function to specify the intrinsic and synaptic properties of neurons exhibit alternative splicing patterns with higher degrees of cell type-specificity compared to differential gene expression ^13^. A prime example is Neurexins (NRXNs), which are a family of synaptic adhesion proteins that play key roles in synaptic organization ^4,14^. Localized to the presynaptic membrane, NRXN isoforms interact with various postsynaptic binding partners such as Neuroligins (NLGNs) and leucine-rich-repeat transmembrane neuronal proteins (LRRTMs) to modulate synaptic properties across the synaptic cleft ^11,15^. In addition to the three *Nrxn* genes (*Nrxn1, Nrxn2,* and *Nrxn3)*, through the use of alternative promoters (*ɑ* and *β* in all three *Nrxn* genes, and an additional *γ* in *Nrxn1*) and alternative splice sites (AS1-AS6), *Nrxn* has the potential to generate thousands of transcript isoforms that when translated, recruit various postsynaptic binding partners ^16,17^. Further, *Nrxn* isoforms exhibit differential expression across select neuron types in the brain and form distinct receptor-ligand complexes, which contribute to specifying cell type-specific synaptic properties ^18–20^. However, the mechanisms that drive cell type-specific *Nrxn* isoform choices remain poorly understood.

Previous studies that investigate *Nrxn* isoform expression have relied on genetic modification to primary transcripts or alternative splice sites. These germline deletion approaches are susceptible to pleiotropic, off-target effects and thus have yielded a wide range of phenotypes that vary with the cellular context. Importantly, these investigations fail to account for potential gain-of-function effects due to shifts in repertoires of *Nrxn* isoforms when interpreting phenotypic outcomes ^21–28^. To address these limitations, we developed a CRISPR epi-editing system in neurons to enable the precise perturbation of select *Nrxn1* isoforms in defined cell types, allowing us to observe downstream regulatory effects. Using this system, we identified transcriptional interference, as well as crosstalk between alternative promoters and splice sites as mechanisms that regulate *Nrxn1* isoform expression in primary neurons. Furthermore, by targeting our epi-editing system to select cell types *in vivo*, we demonstrate that these mechanisms represent a universal principle governing the cell type-specific regulation of *Nrxn1* isoforms in the hippocampus.

## Results and Discussion

### Transcriptional interference revealed by CRISPR epi-editing of *Nrxn1* alternative promoters *in vitro*

To screen gRNAs for CRISPR epi-editing of *Nrxn1*, we designed several gRNAs targeting each of the three *Nrxn1* alternative promoters (*Nrxn1ɑ, Nrxn1β, and Nrxn1γ*; Figure 1A). To confirm the targeting efficiencies of each gRNA, we generated a mouse neuroblastoma cell line (Neuro2A) stably expressing Cas9 via lentivirus, followed by fluorescence-activated cell sorting (FACS) (Figures S1A and S1B). gRNAs were transfected into N2A-Cas9 cells and targeting efficiency was estimated by insertion and deletion (InDel) analysis (Figures S1C and S1D). gRNAs with the highest InDel% for each alternative promoter were selected for epi-editing and packaged in an AAV9 vector with a mCherry fluorescent reporter for neuronal transduction (Figure 1B). Finally, we crossed transgenic floxed CRISPRi mice with CMV-Cre mice to endogenously express dCas9-KRAB across the brain, thereby bypassing the need for delivering the epi-editor, herein referred to as CRISPRi KI mice (Figure 1B) ^29^.

**Figure 1.**
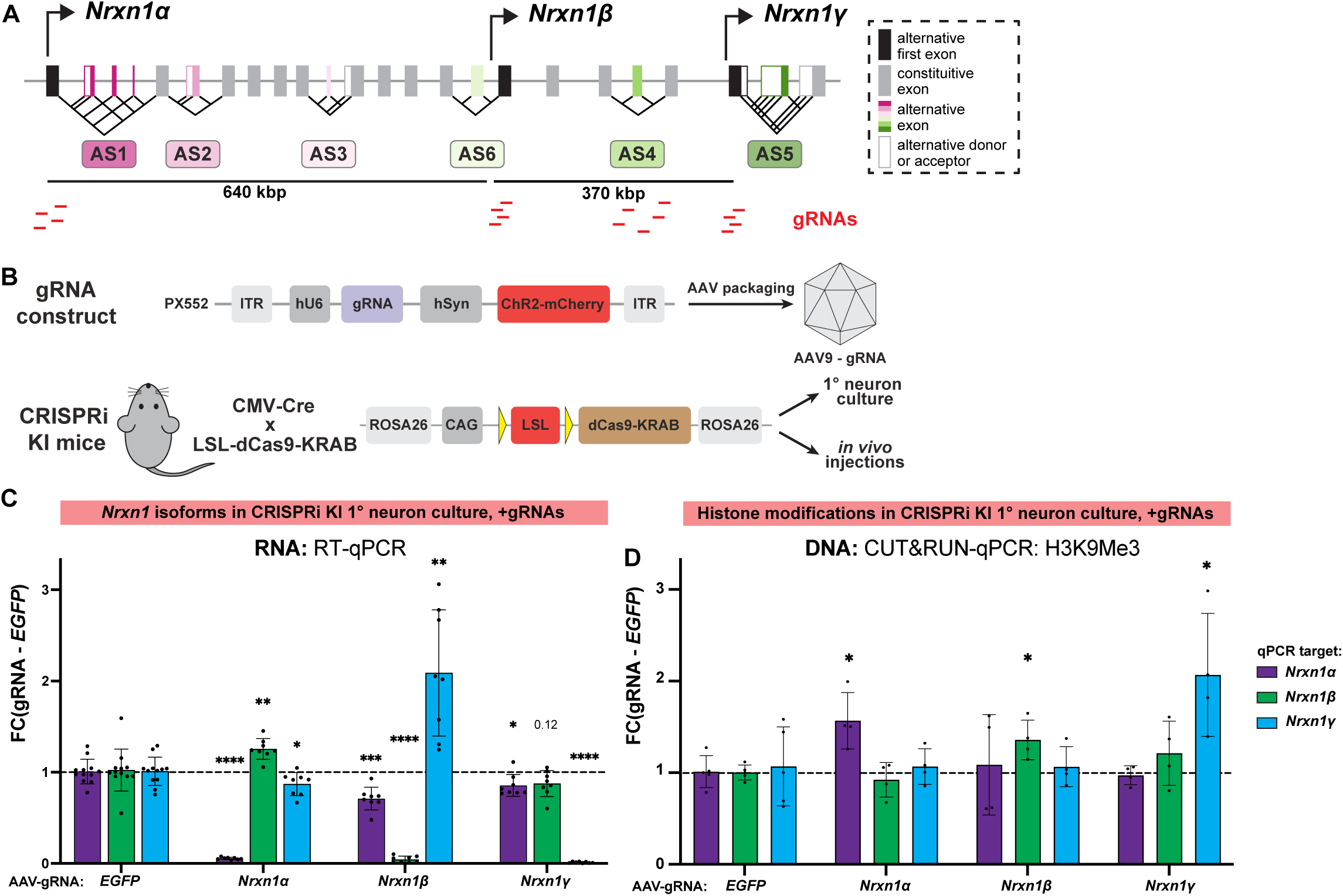
Transcriptional interference revealed by silencing of *Nrxn1* alternative promoters *in vitro*. **(A)** Schematic of the mouse *Nrxn1* gene, marked by the three alternative promoters (α, β, γ) and six alternative splice sites (AS1-AS6). Alternative exons are illustrated in color, constitutive exons in grey, and alternative donors or acceptors in white. Locations of gRNAs are indicated below the gene. Distances not drawn to scale. **(B)** Schematic of the AAV9 construct used for delivery of the gRNAs, and the CRISPRi KI mice generated by crossing CMV-Cre with LSL-dCas9-KRAB. **(C)** RT-qPCR for *Nrxn1* primary transcripts (*α, β, γ*) in primary neurons generated from CRISPRi-KI mice after treatment with AAV-gRNAs (*EGFP* gRNA, *Nrxn1ɑ* gRNA2, *Nrxn1β* gRNA2, and *Nrxn1γ* gRNA2). Expressions of *Nrxn1* primary transcripts were normalized to the expression of *Gapdh*. Fold change (FC) for each gRNA was calculated relative to a non-targeting *EGFP* gRNA. N=8-12 wells per group (* p < 0.05, ** p < 0.01, *** p < 0.001, **** p < 0.0001). *p* values are for gRNAs versus *EGFP* gRNA by unpaired student’s t test. **(D)** CUT&RUN-qPCR against H3K9Me3 for histone modifications around *Nrxn1* alternative promoters in primary neurons generated from CRISPRi-KI mice after treatment with AAV-gRNAs. Enrichments of histone modifications around *Nrxn1* alternative promoters were normalized to histone modifications around the *Gapdh* promoter. Fold change (FC) for each gRNA was calculated relative to a non-targeting *EGFP* gRNA. N=4-5 wells per group (* p < 0.05). *p* values are for gRNAs versus *EGFP* gRNA by unpaired student’s t test.

To measure the efficacy of CRISPRi silencing of *Nrxn1* alternative promoter activity, we introduced promoter-targeting gRNAs to primary cortical and hippocampal neurons prepared from CRISPRi-KI mice. At day *in vitro* 3 (DIV3), AAV9s packaged with gRNA targeting one of the three *Nrxn1* alternative promoters were transduced into the primary neurons, using an *EGFP*-targeting gRNA as a control. At DIV10, one week after AAV transduction, we observed nearly complete silencing (>95%) in the expression of the *Nrxn1* primary transcripts by their respective gRNAs (Figure 1C). Intriguingly, we observed a consistent increase in expression from the downstream promoter of the silenced primary transcript. Namely, *Nrxn1ɑ* silencing increased the expression of the downstream *Nrxn1β* isoform (Figure 1C). Similarly, *Nrxn1β* silencing increased the expression of *Nrxn1γ*.

CRISPRi utilizes dCas9-KRAB to deposit the repressive histone mark H3K9me3 within 500 bps of the transcription start site (TSS) to silence gene expression ^30–32^. To confirm the selectivity of CRISPR epi-editing on target *Nrxn1* promoters, we measured the enrichment of histone modifications with cleavage under targets and release using nuclease (CUT&RUN). In accordance with *Nrxn1* promoter silencing, we observed selective enrichment of H3K9me3-specific heterochromatin regions on the target *ɑ, β, and γ* loci adjacent to their respective gRNAs targets but not on upstream or downstream *Nrxn* promoters (Figure 1D), demonstrating the specificity of our epi-edits. Similarly, chromatin markers of transcriptional activation and elongation, assessing H3K4me3 or H3K36me3 histone modifications, respectively, did not reveal chromatin remodeling at the non-target alternative promoters (Figure S2). Since H3K4me3 marks active promoters, we hypothesized that increases or decreases in the expression levels of *Nrxn1* primary transcripts would be directly linked to H3K4me3 levels. However, we observed weak correlations between the expression of primary transcript levels with H3K4me3-associated chromatin marks (Figures 1C and S2B). This suggests that the enrichment of the repressive H3K9me3 histone mark was sufficient for gene silencing at gRNA-targeted promoters (Figure 1D) ^33^. Our data further demonstrates that gene activation at downstream *Nrxn1* promoters occurred independently of H3K4me3 enrichment. Given the specificity of CRISPRi in targeting the individual *Nrxn1* promoters, we hypothesized that the relationship between upstream silencing and downstream activation of *Nrxn1* primary transcripts reflects the intrinsic relationship between alternative promoter usage in *Nrxn1*.

Transcriptional interference is a phenomenon in which the activity of one promoter can modulate the expression of a neighboring promoter in *cis*, often due to the passage of RNA polymerase II (Pol II) initiated from the upstream site ^34–37^. Naturally occurring cases of transcriptional interference have been observed between promoters separated by hundreds of base pairs, typically in bacteria or yeast. ^38–42^. Here, we describe a striking example of long-range transcriptional interference in the mammalian brain involving *Nrxn1* alternative promoters that are separated by hundreds of kilobases. Specifically, the *Nrxn1ɑ* and *Nrxn1β* promoters are located ∼640,000 bps apart, while the *Nrxn1β* and *Nrxn1γ* are ∼370,000 bps apart (Figure 1A). We propose that transcription initiated from an upstream alternative promoter interferes with downstream promoter activity by transiently occupying the downstream site with elongating RNA pol II. By silencing the upstream promoter, we relieve this interference, allowing transcription to initiate at the downstream promoter. For example, silencing the Nrxn1α promoter permits increased transcription from the Nrxn1β promoter, and silencing Nrxn1β enables activation of Nrxn1γ.

Although transcriptional interference is described for individual genes arranged in tandem and involved in regulatory feedback loops, our findings reveal a unique instance in which this mechanism operates between alternative promoters within a single gene that shares most of the same open reading frame. This arrangement enables *cis-*regulatory control within a single gene to generate distinct isoforms that support diverse synaptic functions in neurons. We further hypothesize that this mode of isoform regulation may be widespread and detectable using epi-editing approaches, as suggested by a recent study in which CRISPRi unexpectedly activated a nested gene within an intron of its target gene, likely through a similar transcriptional interference mechanism ^43^.

We also noted that silencing the *Nrxn1β* promoter modestly decreased the expression of *Nrxn1ɑ* in the upstream direction, but to a smaller extent than with transcription interference (Figure 2B). Structural evidence demonstrates that CRISPR-based targeting within a gene body influences transcription in a strand-specific manner due to dCas9-associated gRNA annealing to the non-template strand, thus slowing down RNA pol II by impeding the progression of the elongation complex ^43,44^. The recruitment of dCas9-KRAB to the *Nrxn1β* promoter also falls within the gene body of *Nrxn1ɑ*. Thus, it is likely that *Nrxn1β* silencing reduced *Nrxn1ɑ* expression due to dCas9-KRAB functioning as a roadblock for RNA pol II elongation across the *Nrxn1ɑ* gene body.

**Figure 2.**
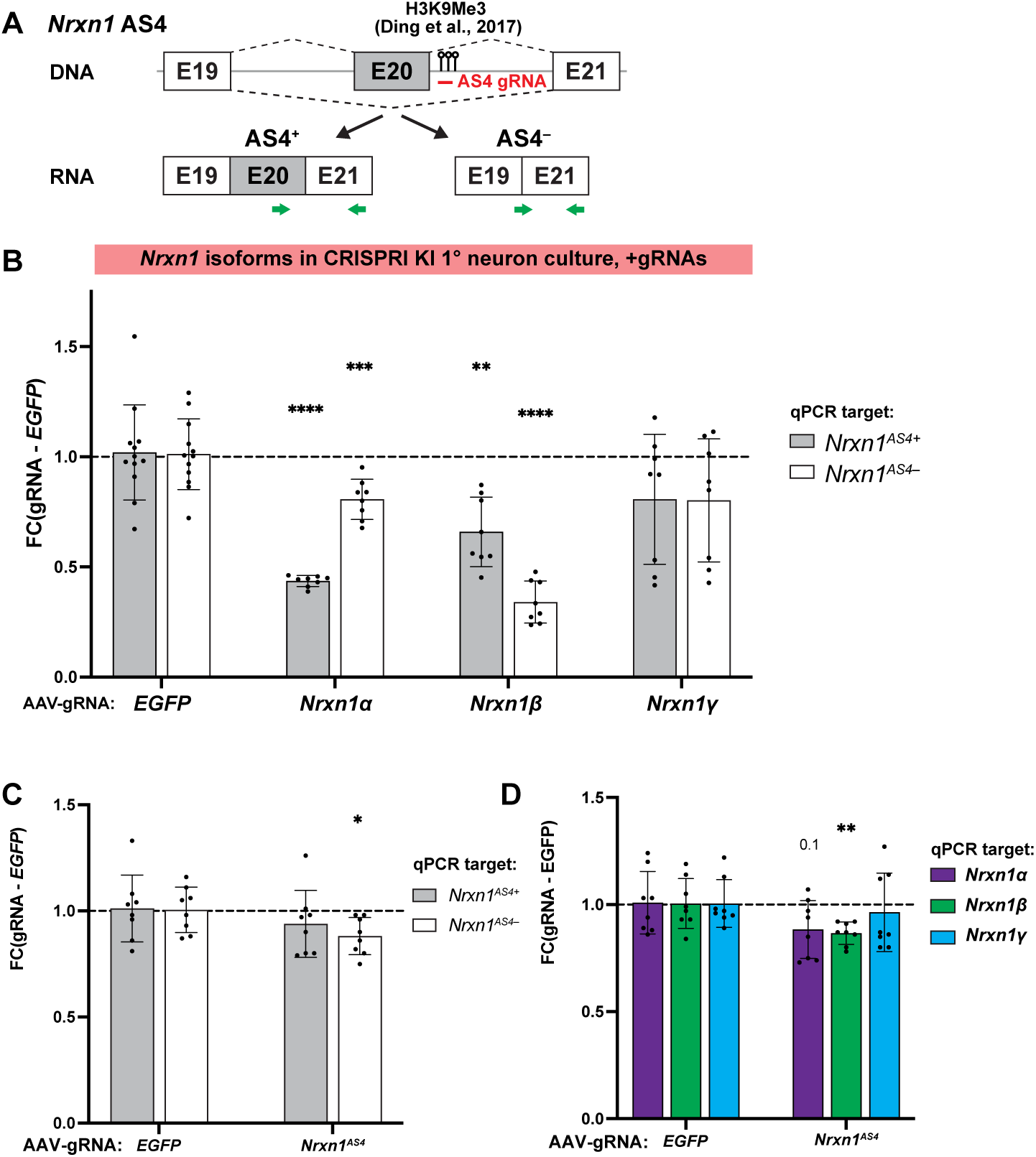
Splicing at AS4 is affected by alternative promoter usage, but not by intragenic CRISPR epi-editing. **(A)** Schematic of *Nrxn1* AS4 splicing. RT-qPCR primers for quantifying AS4^+^ and AS4^−^ splicing. AS4-gRNA1 for intragenic CRISPR epi-editing targets the genomic region where H3K9Me3 enrichment was identified in previously published study. **(B)** RT-qPCR for *Nrxn1* AS4 splicing in primary neurons generated from CRISPRi-KI mice after treatment with AAV-gRNAs targeting the individual alternative promoters. Expressions of *Nrxn1* AS4+ or AS4-were normalized to the expression of *Gapdh*. FC for each gRNA was calculated relative to a non-targeting *EGFP* gRNA. N=8-12 wells per group (** p < 0.01, *** p < 0.001, **** p < 0.0001). *p* values are for gRNAs versus *EGFP* gRNA by unpaired student’s t test. **(C-D)** RT-qPCR for **(C)** *Nrxn1* AS4 splicing and **(D)** *Nrxn1* primary transcripts after treatment with AAV-AS4-gRNA1. Expressions of *Nrxn1* transcripts were normalized to the expression of *Gapdh*. Fold change (FC) for AS4-gRNA1 was calculated relative to a non-targeting *EGFP* gRNA. N=8 wells per group (* p < 0.05, ** p < 0.01). *p* values are for AS4-gRNA1 versus *EGFP* gRNA by unpaired student’s t test.

### Splicing at *Nrxn1* AS4 is guided by alternative promoter usage, not by intragenic CRISPR epi-editing

Both *Nrxn1ɑ* and *Nrxn1β* primary transcripts undergo alternative splicing at AS4, while transcription from the *Nrxn1γ* promoter occurs downstream of AS4 (Figure 1A). Inclusion or exclusion of the alternative exon 20 generates an AS4^+^ and AS4^−^ isoform, respectively (Figure 2A). The functional consequence of splicing at *Nrxn1* AS4 modifies the ligand-binding surface of NRXN1, altering its affinity to post-synaptic binding partners to modulate synaptic strength ^27^. When silencing either the Nrxn1α or Nrxn1β alternative promoters in primary neurons with CRISPRi, we observed a striking, disproportionate effect on downstream AS4 splicing (Figure 2B). Specifically, silencing Nrxn1α led to a marked reduction in AS4^+^ isoforms, while silencing Nrxn1β resulted in a significant decrease in AS4^-^ isoforms. These results indicate a strong link between alternative promoter usage and AS4 splice site selection, with Nrxn1α transcripts being predominantly AS4^+^, whereas Nrxn1β transcripts are mainly AS4^-^ (Figures 1C and 2B). The potential for *cis*-regulatory crosstalk between the *Nrxn1ɑ* or *Nrxn1β* promoter with the AS4 splice site, independent of trans-regulatory splicing factors, is supported by previous studies demonstrating effects of promoter usage on splicing via promoter swapping ^45^. For example, differences in promoter architecture can recruit factors that influence the RNA pol II elongation rate and the spatiotemporal dynamics of spliceosome assembly and is an attractive mechanism for influencing splicing decisions ^46–48^.

Previous studies suggest that activity-dependent enrichment of repressive H3K9me3 marks at *Nrxn1* AS4 influences the kinetics of RNA pol II and regulates splicing ^49^. Thus, we employed intragenic CRISPR epi-editing to assess how local epigenetic modifications influence alternative splicing at *Nrxn1* AS4. We hypothesize that depositing H3K9me3 via dCas9-KRAB adjacent to AS4 could slow RNA pol II elongation and recapitulate previously observed increases in *Nrxn1* AS4 inclusion ^49^. Using our N2A system, we identified a gRNA that targeted the AS4 region previously associated with H3K9Me3 deposition and RNA pol II pausing (Figures 2A and S1D) ^49^. Following AAV delivery of AS4-targeting gRNA to primary neurons, contrary to expectations, targeted epi-editing at AS4 did not affect splice inclusion, with modest decreases in *Nrxn1* AS4^−^ (Figure 2C). We hypothesized that epi-editing at the downstream *Nrxn1* AS4 locus may influence upstream promoter usage. Thus, we measured primary *Nrxn1* transcript levels and observed an associated decrease in *Nrxn1β* expression (Figure 2D). Given our finding that *Nrxn1β* isoforms are predominantly AS4^−^, local epi-editing at the AS4 locus likely decreases the expression of the *Nrxn1β* primary transcript rather than influencing AS4 splicing decisions. Our data suggest that AS4 splicing is not regulated by RNA pol II kinetics, indicating that neither promoter-recruited factors nor local intragenic histone modifications significantly influence AS4 splicing by changing RNA pol II rate.

Previous studies that implicated the role of H3K9me3 on AS4 splicing relied on *Suv39h1-* deficiency, a histone methyltransferase that regulates H3K9me3 levels ^49^. However, H3K9me3 is critical for regulating gene transcription at promoters on a genome-wide level. Given our observation that AS4 splicing is closely linked to alternative promoter usage, a global change that alters *Nrxn1* primary transcript levels would also shift levels of AS4 isoforms. To disentangle the potential roles of alternative promoter choice on AS4 splicing, our targeted CRISPR epi-editing approach distinguishes direct roles of local H3K9me3 on splicing from the indirect roles on gene expression. However, although we did not observe changes in *Nrxn1* AS4 inclusion with intragenic epi-editing using dCas9-KRAB, epi-editors that deposit other histone modifications may still have effects in different genomic contexts ^50^. Instead, we hypothesize that splicing factors directly recruited to promoters and brought to splice sites by the C-terminal domain of RNA pol II may play a role in influencing AS4 splicing decisions ^51^. The KH-domain RNA binding protein SLM2 has been shown to regulate splicing at *Nrxn* AS4 ^52^. To further investigate this promoter-splicing relationship, we next examined this in an *in vivo* context in the mouse hippocampus, where SLM2 expression and its effect on AS4 splicing is well-defined ^18^. In parallel, we explore the promoter-promoter relationship, as *Nrxn1ɑ* and *Nrxn1β* promoter usage differs in select cell types of the hippocampus ^27,53^.

### *In vivo* epi-editing of *Nrxn1* alternative promoters in the mouse hippocampus

To determine whether transcriptional interference of *Nrxn1* alternative promoters occurs *in vivo*, we applied our CRISPR epi-editing system to the mouse hippocampus. We recapitulated the cell type-specific expression patterns of *Nrxn1* primary transcripts previously identified by *in situ* hybridization ^18^, using RT-qPCR on micro-dissected CA1 and CA3. We confirmed high levels of *Nrxn1ɑ* and minimal to no *Nrxn1β* in CA1, with the inverse expression pattern in CA3, namely low levels of *Nrxn1ɑ* and high levels of *Nrxn1β* (Figure 3A). Consistent with our hypothesis that *Nrxn1* promoter choice is coupled to downstream AS4 decisions, we observed that the elevated expression of *Nrxn1ɑ* in CA1 was coupled with higher levels of AS4^+^ (Figure 3A), whereas higher expression of *Nrxn1β* in CA3 corresponded with higher levels of AS4^−^ isoforms. We observed no significant differences in *Nrxn1γ* expression between CA1 and CA3.

**Figure 3.**
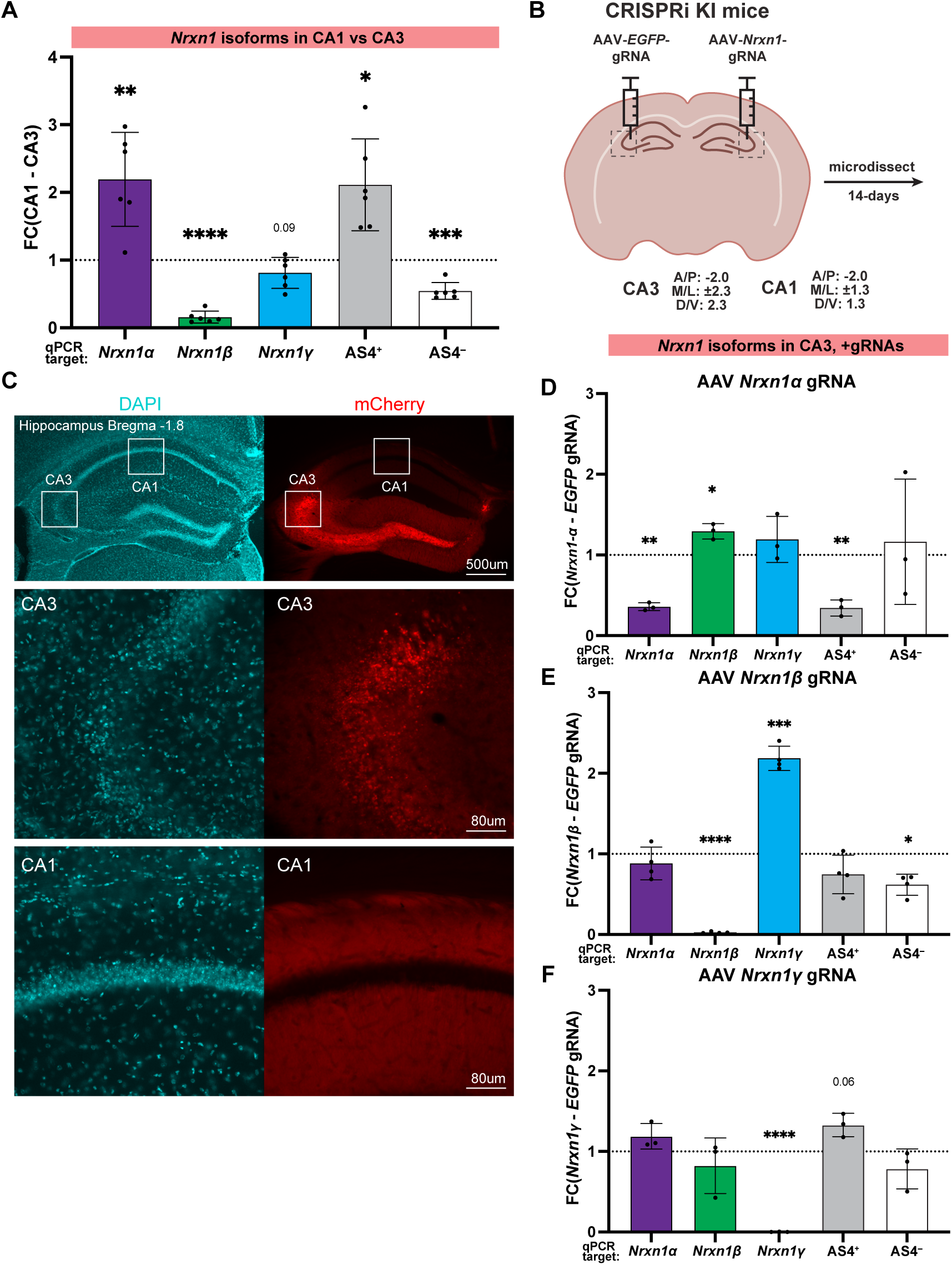
*In vivo* epi-editing of *Nrxn1* alternative promoters in CA3 of the mouse hippocampus. **(A)** RT-qPCR for *Nrxn1* transcripts in the WT mouse hippocampus. Expressions of *Nrxn1* transcripts were normalized to the expression of *Gapdh*. FC were calculated by comparing expressions in CA1 versus CA3. N=6 hippocampi (* p < 0.05, ** p < 0.01, *** p < 0.001, **** p < 0.0001). *p* values from one-sample t test. **(B)** Schematic for bilateral stereotaxic injections of AAV-gRNAs into the hippocampus of CRISPRi KI mouse, with *EGFP* gRNA and *Nrxn1* alternative promoters gRNA to the contralateral and ipsilateral hemispheres, respectively. Tissues were harvested 14-days post injection. Stereotaxic coordinates for the injections are indicated. **(C)** Representative images of DAPI and mCherry fluorescence in the hippocampus 14-days post injection to CA3. Insets for CA3 and CA1 are shown. Scale bars are 500 µm (top) and 80 µm (bottom). **(D-F)** RT-qPCR for *Nrxn1* transcripts in CA3 after injection of **(D)** *Nrxn1ɑ*-gRNA, **(E)** *Nrxn1β*-gRNA, and **(F)** *Nrxn1γ*-gRNA. Expressions of *Nrxn1* transcripts were normalized to the expression of *Gapdh*. FC were calculated by comparing expressions in the *Nrxn1* gRNA-injected hemisphere against the *EGFP* gRNA-injected hemisphere within the same animal. N=3-4 mice per group (* p < 0.05, ** p < 0.01, *** p < 0.001, **** p < 0.0001). *p* values from one-sample t test.

To assess the effect of epi-editing on *Nrxn1* promoter-promoter relationships *in vivo*, we stereotactically injected AAVs with gRNAs targeting each *Nrxn1* alternative promoter to CRISPRi-KI mice. We first targeted the CA3 region, as CA3 pyramidal neurons express all three *Nrxn1* primary transcripts (Figure 3B). We confirmed on-target AAV delivery to CA3 by selective mCherry expression in CA3 pyramidal neurons and their axonal projections to CA1, with no detectable mCherry in the pyramidal layer of non-targeted CA1 (Figure 3C). Consistent with our primary neuron data, we robustly silenced the *Nrxn1ɑ, Nrxn1β, and Nrxn1γ* alternative promoters with their respective gRNAs (Figures 3D-F). Remarkably, we recapitulated the promoter crosstalk observed in primary neuron culture, as the silencing of *Nrxn1ɑ* and *Nrxn1β* in CA3 also increased expression of *Nrxn1β* and *Nrxn1γ*, respectively (Figures 1C and 3D-F). Similarly, *Nrxn1ɑ* and *Nrxn1β* silencing disproportionately altered AS4^+^ and AS4^−^ isoforms, confirming that downstream AS4 choice is coupled to promoter choice *in vivo* (Figure 2B and 3D-F).

To confirm that transcriptional interference is sufficient to drive alternative promoter usage of *Nrxn* isoforms in the hippocampus, we targeted the CA1 region where *Nrxn1β* expression is absent (Figure 3A). We hypothesize that low *Nrxn1β* levels in CA1 arise from robust engagement of the *Nrxn1α* promoter driving transcriptional interference at the *Nrxnβ* promoter, and that relieving this interference would activate *Nrxn1β* expression in CA1 pyramidal neurons. By targeting *Nrxn1α* with a gRNA and confirming on-target AAV delivery to CA1 via mCherry expression (Figure 4A), we found that strong silencing of *Nrxn1α* primary transcripts increased *Nrxn1β* expression in CA1 (Figure 4B). Together, our results show that transcriptional interference is not only necessary for regulating promoter usage in CA3 but that by removing this interference alone is sufficient to activate otherwise repressed isoforms in CA1 without the need for additional transcription factors to activate *Nrxn1β* expression.

**Figure 4.**
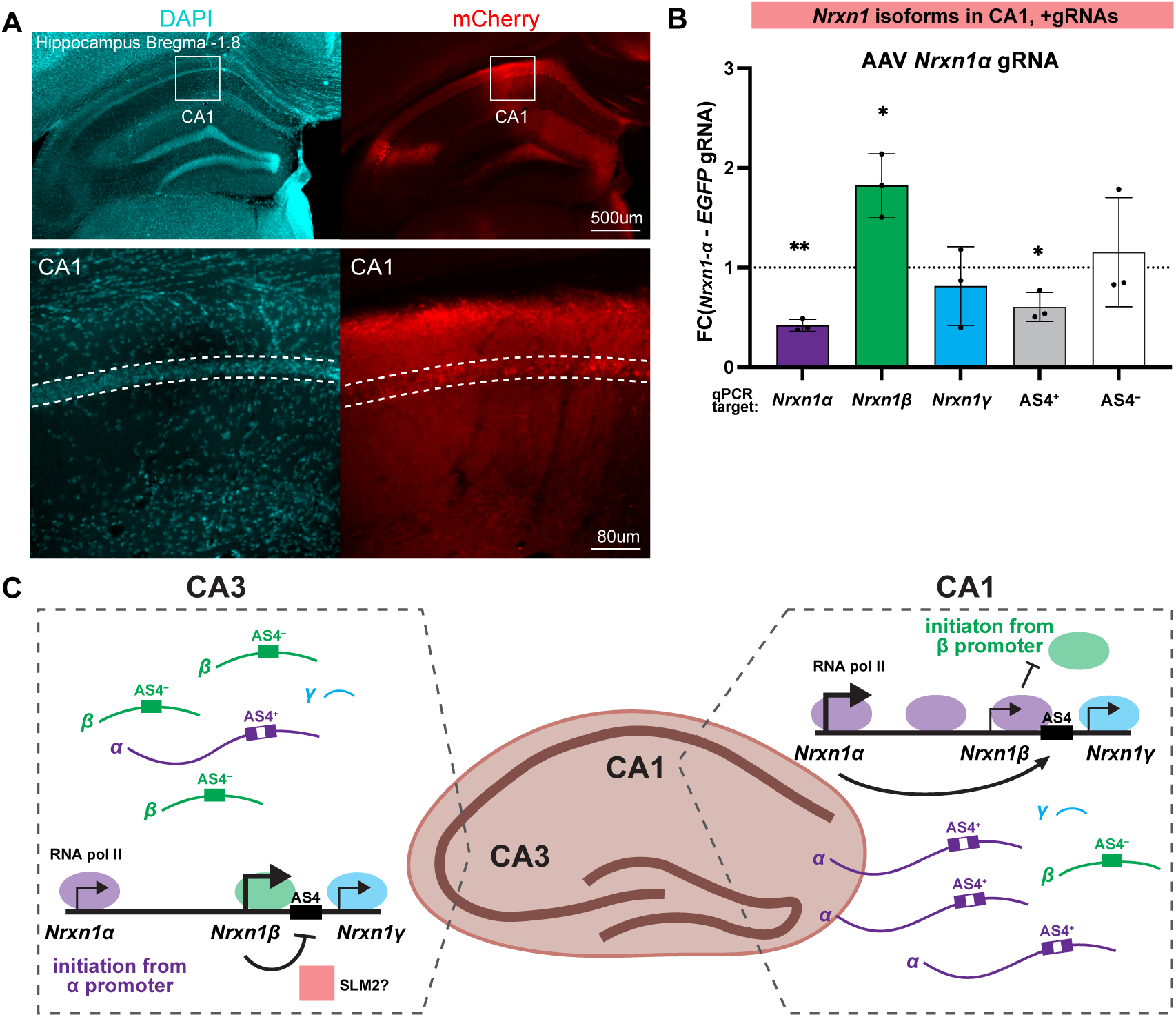
Transcriptional interference dictates cell type-specific expression of Nrxn1 isoforms across the hippocampus. **(A)** Representative images of DAPI and mCherry fluorescence in the hippocampus 14-days post injection into CA1. Inset for CA1 is shown. Scale bars are 500 µm (top) and 80 µm (bottom). **(B)** RT-qPCR for *Nrxn1* transcripts in CA1 after injection of *Nrxn1ɑ*-gRNA. Expressions of *Nrxn1* transcripts were normalized to the expression of *Gapdh*. FC were calculated by comparing expressions in the *Nrxn1* gRNA-injected hemisphere against the *EGFP* gRNA-injected hemisphere within the same animal. N=3 mice per group (* p < 0.05, ** p < 0.01). *p* values from one-sample t test. **(C)** Schematic of the cell-type specific regulation of *Nrxn1* isoforms in the hippocampus across alternative promoters and splice sites via transcriptional interference.

Together, we discovered that transcriptional interference is a key mechanism that regulates alternative promoter usage of the synaptic organizer, *Nrxn1.* We found that altering the expression of *Nrxn1ɑ* directly tunes levels of *Nrxn1β.* We demonstrate that transcriptional interference is necessary and sufficient to regulate patterns of *Nrxn1* primary transcripts in the mouse hippocampus by inversely controlling *Nrxn1ɑ* levels in CA1 pyramidal neurons and *Nrxn1β* levels in CA3 pyramidal neurons (Figures 4C). Since the larger NRXNɑ protein recruits distinct macromolecular assemblies across the synaptic cleft compared to the shorter NRXNβ ^11^, our results reveal that transcriptional interference is a key mechanism for governing distinct patterns of synaptic connectivity. Surprisingly, we showed that *Nrxn1ɑ* and *Nrxn1β* isoforms exhibit bias towards AS4^+^ and AS4^−^, respectively. Previous studies showed the generation of *Nrxn1 AS4^+^* or *Nrxn1 AS4^-^* isoforms to be controlled by the splicing factor SLM2, which exhibits differential expression between principal neurons and interneurons in the hippocampus ^18^. However, SLM2 expression is comparable across the principal neurons in the individual hippocampal subfields ^54^. Thus, differences in SLM2 expression cannot explain the differences in AS4 splicing between CA1 and CA3 pyramidal neurons. Here, we postulate that the coupling of alternative splicing to transcription provides an additional layer of regulation - independent of local chromatin modifications - where promoter choice alone can dictate splicing decisions without the need for differential expression of splicing factors (Figure 4C). In summary, we developed a platform for CRISPR epi-editing in neurons to reveal the regulatory relationships between alternative promoters and alternative splicing and identify transcriptional interference as a critical mechanism that generates heterogeneity for cell type-specific synaptic functions in the brain.

## Resource Availability

### Lead contact

Further information and requests for resources should be directed to Dr. Andrea Gomez.

### Materials availability

This study did not generate new unique reagents or materials.

### Data and code availability

Any additional information required to reanalyze the data reported in this paper is available from the lead contact upon request.

## Acknowledgements

We thank J. Nuñez and F. Urnov for the N2A cells and plasmids used in the N2A-Cas9 experiments. Y.H. was financially supported by a National Institutes of Health T32 grant (5T32GM007232) and a Predoctoral Fellowship from the PhRMA Foundation. This work was supported by funds to A.M.G. by a NARSAD Young Investigator Grant from the Brain & Behavior Research Foundation, the Shurl and Kay Curci Foundation, the Rennie Fund for the Study of Epilepsy, and a IDOR-Innovative Genomics Institute grant. Sanger sequencing was performed by the UC Berkeley DNA Sequencing Facility. FACS was conducted at the Flow Cytometry Core Facility at UC Berkeley.

## Author contributions

Y.H. and A.M.G. conceived and designed the experiments. Y.H. conducted all experiments. Y.H., and A.M.G. wrote the manuscript. Both authors reviewed the manuscript before submission.

## Supplemental figure titles and legends

**Supplemental Figure 1. *in vitro* platform to screen targeting efficiencies of gRNAs in neurons**

**(A)** Schematic of CRISPR constructs used to generate N2A-Cas9 and deliver the gRNAs, as well as pipeline for assessing gRNA targeting efficiencies in N2A-Cas9.

**(B)** FACS for BFP expression in WT N2A, N2A 48-hours post infection with pMH004, and N2A-Cas9 after having been grown out. Percentage of BFP+ cells are indicated.

**(C)** Representative traces of gRNA targeting efficiencies for *Nrxn1ɑ* gRNA2 against pLG1 non-targeting (NT) gRNA calculated by ICE analysis. *Nrxn1ɑ* gRNA2 sequence is indicated by a solid black line, the protospacer adjacent motif (PAM) is indicated by a dashed red line, and the Cas9 cut site is indicated by a dashed black line.

**(D)** Quantification of targeting efficiencies for gRNAs against the three *Nrxn1* alternative promoters and AS4 splice site in N2A-Cas9 by InDels.

**Supplemental Figure 2. Epigenetics of transcriptional interference at *Nrxn1* alternative promoters**

**(A)** Schematic of primers used to examine enrichment of histone modifications at individual alternative promoters by CUT&RUN-qPCR.

**(B-C)** CUT&RUN-qPCR against **(B)** H3K4Me3 and (**C)** H3K36Me3 for histone modifications around *Nrxn1* alternative promoters in primary neurons generated from CRISPRi-KI mice after treatment with AAV-gRNAs. Enrichments of histone modifications around *Nrxn1* alternative promoters were normalized to histone modifications around the *Gapdh* promoter. FC for each gRNA was calculated relative to a non-targeting *EGFP* gRNA. N=3-5 wells per group (* p < 0.05). *p* values are for gRNAs versus *EGFP* gRNA by unpaired student’s t test.

## Method details

### Mice

dCas9-KRAB mice and CMV-Cre mice were obtained from Jackson Laboratories (Jax stock no: 033066, 006054). All lines were maintained on a C57Bl6/J background. dCas9-KRAB mice were crossed with CMV-Cre mice to generate knock-in mice with ubiquitous expression of the dCas9-KRAB fusion protein. All experiments were performed in accordance with the regulations of the Institutional Animal Care and Use Committee (IACUC) of the University of California, Berkeley.

### Generation of Neuro2A-Cas9-expressing stable cell line

The Cas9 lentivirus plasmid, pMH004 was a gift from Jonathan Weissman (Addgene plasmid # 174087; http://n2t.net/addgene:174087; RRID:Addgene_174087). Lentiviral particles were produced by co-transfection of HEK293T cells with 1.5μg of Cas9 plasmid and standard packaging vectors using Mirus TransIT-LT1 Transfection Reagent (Mirus MIR2300). Media was changed 24 hours post-transfection with DMEM+++ (DMEM (Sigma Aldrich D5796) containing 10% fetal bovine serum and 1% penicillin/streptomycin). Viral supernatant was collected 48 hours post-transfection and filtered through a 0.45 mm syringe filter. N2A cells were cultured at 37°C and 5% CO_2_ in DMEM+++. For lentiviral transduction, N2A cells were seeded in 6-well plates and immediately transduced with 150μl of lentivirus. Two days following infection, cells were sorted with FACS and BFP+ cells collected for downstream experiments.

### Targeting efficiency of gRNAs in N2A-Cas9

The gRNA plasmid, pLG1 (pU6-sgRNA Ef1alpha-Puro-T2A-BFP) was a gift from Luke Gilbert (Addgene plasmid # 217305; http://n2t.net/addgene:217305; RRID:Addgene_217305), with BFP replaced by GFP. gRNAs were designed using The Broad Institute’s gRNA design tool and cloned in by annealed oligo cloning using BstXI and BlpI restriction enzymes following previously published protocol. For transfection, N2A-Cas9 cells were seeded into 12-well plates and transfected the following day with 1μg of gRNA plasmids using Lipofectamine-2000 (Invitrogen 11668019). Cells were harvested 48 hours post-transfection with QuickExtract DNA extraction solution (Lucigen QE09050) by following manufacturer’s protocol. Amplicons were generated using Promega PCR master mix (Promega M7502) and Sanger sequencing was performed by the UC Berkeley DNA Sequencing Facility. Targeting efficiency was calculated using Synthego’s Inference of CRISPR Edits (ICE), comparing target-specific gRNAs to pLG1 empty vector.

### Primary cell culture

Primary mixed cortical/hippocampal cultures were prepared from P0 mouse pups. Brains were dissected in 1x HBSS (Gibco 14185052) with 10mM HEPES (Gibco 15630080). Cortices and hippocampi were collected and minced before transferring to a 15mL tube with 1mL .05% trypsin-EDTA (Gibco 25300062) and 100μL 1mg/mL DNase I (Roche10104159001) and incubated for 5 mins in a 37°C water bath. Another 100μL of DNase I was added and incubated for an additional 10 mins with occasional agitation. Subsequently, the digestion solution was removed, and the tissue washed twice with DMEM+++ before triturating in 1mL of DMEM+++ with DNase I and soybean trypsin inhibitor (Sigma Aldrich T6522), 8x each in a 1mL pipette tip and a 200μL pipette tip. After trituration, 2mL of DMEM+++ was added and the cell suspension filtered through a 40μm cell strainer before centrifuging at 100g for 10 mins. The supernatant was removed, and the cell pellet was resuspended in DMEM+++. Cell count was determined with a hemocytometer, and 100,000 cells per well were plated in a 24-well plate coated with poly-D-lysine. After 2-4 hours, the DMEM+++ plating media was replaced with NB+++ (Neurobasal media (Gibco 21103049) containing 1x B27 (Gibco 17504044) and 2mM GlutaMAX (Gibco 35050061)).

### Design and cloning of AAV-gRNA

The AAV-gRNA vector, PX552, was a gift from Feng Zhang (Addgene plasmid # 60958; http://n2t.net/addgene:60958; RRID:Addgene_60958), with EGFP replaced by ChR2(H134R)-mCherry. gRNAs were cloned in by annealed oligo cloning using SapI restriction enzyme, following previously published protocol on PX552’s Addgene page.

### AAV production and viral transduction in vitro

AAV9 production was performed by Vector Biolabs. Packaged AAVs ranged in titers from 1 to 2 x 10^13^ GC/mL. AAVs were transduced into primary neurons on DIV3 at an MOI of 120,000 and incubated for 48 hours, before a half-media switch on DIV5.

### RNA extraction from primary cultures, reverse transcription and RT-qPCR

RNA extraction from primary neurons was done on DIV10. Cells were lysed in-well with 350 uL of RLT buffer from RNeasy Mini kit (Qiagen 74106), and RNA was purified by following the manufacturer’s protocol. 100 ng of RNA from each sample was used for reverse transcription with SuperScript III First Strand Synthesis SuperMix (Invitrogen 11752250). RT-qPCR was performed with PowerUp SYBR Green PCR Master Mix (Applied Biosystems 4368706) on a QuantStudio 3. The 2^-ΔΔCT^ method was used to calculate relative expression between *Nrxn1* versus *EGFP* gRNA infected neurons. Statistical analysis was performed using Graph Pad Prism (v.10.3.1). Two-tailed unpaired Student’s t-test was used to compare means.

### CUT&RUN qPCR

CUT&RUN was performed following the manufacturer’s protocol (Cell Signaling Technology 86652). In brief, primary neurons were detached on DIV10 using pre-warmed Accutase (Corning 25058CI) and incubated for 15 mins at 37°C. Wells were flooded with Neurobasal media (Gibco 21103049) and triturated into single cell suspension. Cells were centrifuged at 600g for 3 mins and washed twice with Wash Buffer (1X wash buffer, 1X spermidine, and 1X Protease Inhibitor Cocktail). Concanavalin A magnetic beads were pre-activated by washing twice with Activation Buffer on a magnetic rack and resuspended in Binding Buffer (1X antibody binding buffer, 1X spermidine, 1X Protease Inhibitor Cocktail, 0.025% Digitonin). Cells were incubated with Concanavalin A magnetic beads for 5 mins at room temperature before 2μg of antibodies targeted to specific histone modifications were added for overnight incubation at 4°C (rabbit anti-H3K9Me3 (Active Motif 39062), rabbit anti-H3K4Me3 (Cell Signaling Technology 9751), rabbit anti-H3K36Me3(Active Motif 61902)). Samples were washed twice on a magnetic rack with Digitonin Buffer (1X wash buffer, 1X spermidine, 1X Protease Inhibitor Cocktail, 0.025% Digitonin) before incubating with pAG-MNase for 1 hour at 4°C. Samples were washed twice with Digitonin Buffer again, and left on ice for 5 mins before activation of pAG-MNase with CaCl_2_ for 30 mins at 4°C. Digestion was stopped with Stop Buffer (1X stop buffer, 0.025% Digitonin, 7.5μg of RNase A). Samples were incubated at 37°C for 10 mins and centrifuged at 16,000g for 2 mins at 4°C. DNA fragments were purified with Monarch PCR & DNA Cleanup Kit (New England Biolabs T1030) by following the manufacturer’s protocol and eluted in 10uL water. CUT&RUN qPCR was performed with PowerUp SYBR Green PCR Master Mix on a QuantStudio 3.

### Stereotaxic injections in vivo

Adult mice were anesthetized by inhalation of isofluorane (induction at 4% and maintenance at 2%) delivered with constant oxygen flow (1 L/min). Analgesics meloxicam (5 mg/kg) and lidocaine (2%, up to .025 mL) were injected subcutaneously and locally at the surgical site, respectively, prior to incision. Mice were immobilized on a stereotaxic apparatus (WPI) with ear bars and a tooth bar. After incision, a 1mm hole was drilled on the surface of skull in each hemisphere, and a 36-gauge NanoFil needle was used for injection into the CA3 of the hippocampus (coordinates: A/P: −2.0, M/L: +/- 2.3, D/V: −2.3). 1uL of AAV-gRNA was injected into each hemisphere such that each animal had a control and experimental side, at 383 nL/min with a microinjection syringe pump (WPI UMP3T). After injection, the needle was left in for 2 mins to prevent backflow before it was slowly raised out of the brain. The surgical site was then closed with 4-0 Visorb sutures (Stoelting). Meloxicam (5 mg/kg) was again injected subcutaneously immediately after surgery. Animals were allowed to recover in a warmed cage on a heat pad before returning to their home cages to be monitored for the next 7 days.

### RNA extraction from surgical animals

Animals were sacrificed 14 days after surgery with 4% isofluorane and transcardially perfused with ice cold PBS. 300um slices were cut coronally using a vibratome (Leica Microsystems VT1200S). Individual slices were then microdissected based on mCherry fluorescence to isolate the CA3 region, +/- 500um anterior/posterior to the injection site. Tissues were lysed with 350 uL of RLT buffer from RNeasy Mini kit by triturating 8x in a 1mL pipette tip and 8x in a 200uL pipette tip before centrifuging for 3 mins at 18,000 x g. RNA was then purified by following the manufacturer’s protocol. RT-qPCR was performed as described previously. The 2^-ΔΔCT^ method was used to calculate relative expression between *Nrxn1* versus *EGFP* gRNA injected hemispheres within each animal. One-sample t-test was used to determine if means differ from 1.

## Supplemental Information

Supplemental Figures 1-2.

