## Supplemental Figures for "CRISPR epi-editing reveals regulatory crosstalk between alternative promoters and splicing in Neurexin isoform expression"

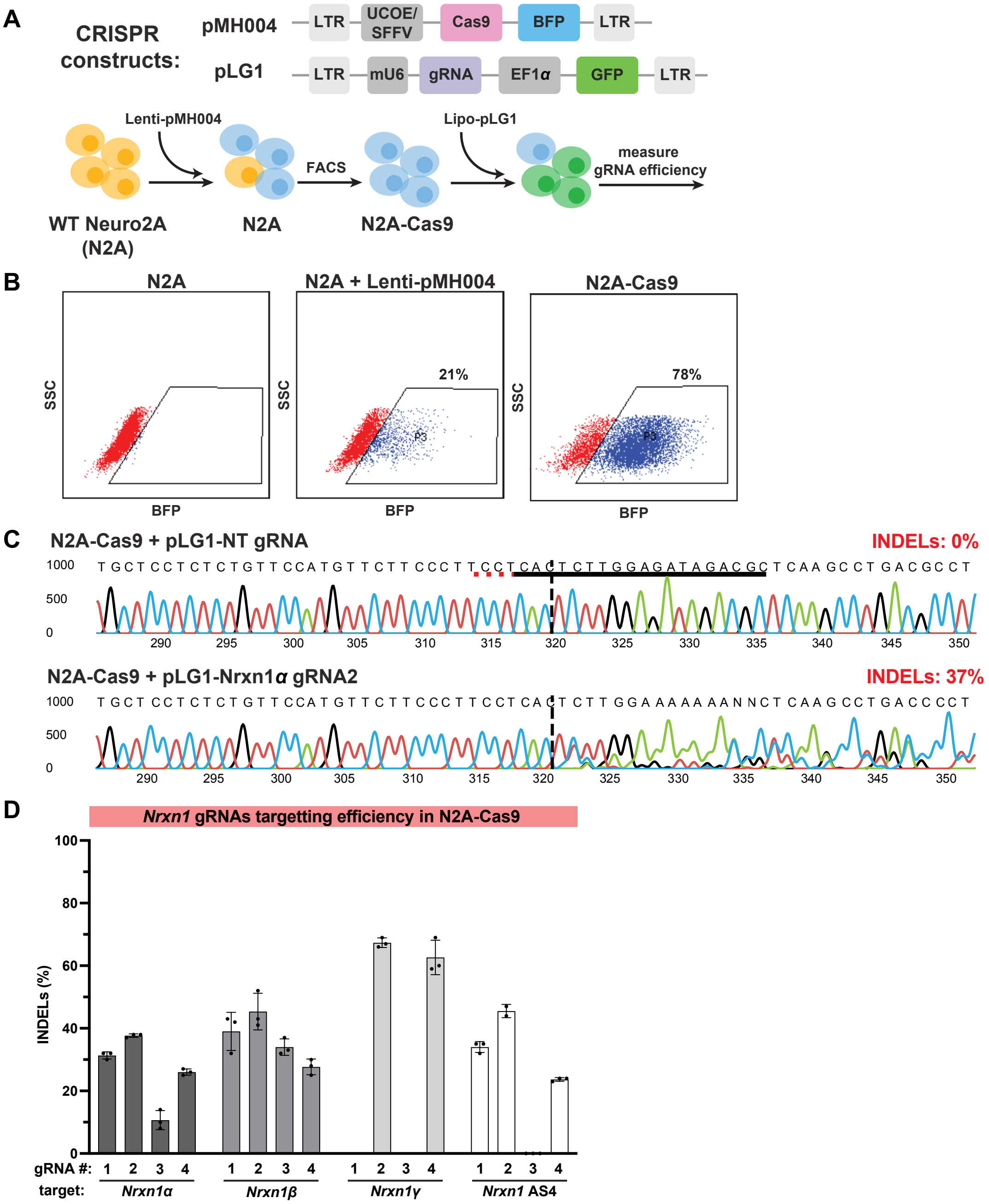


**Supplemental Figure 1. in vitro platform to screen targeting efficiencies of gRNAs in neurons**

(A) Schematic of CRISPR constructs used to generate N2A-Cas9 and deliver the gRNAs, as well as pipeline for assessing gRNA targeting efficiencies in N2A-Cas9.

(B) FACS for BFP expression in WT N2A, N2A 48-hours post infection with pMH004, and N2A-Cas9 after having been grown out. Percentage of BFP+ cells are indicated.

(C) Representative traces of gRNA targeting efficiencies for Nrxn1α gRNA2 against pLG1 non-targeting (NT) gRNA calculated by ICE analysis. Nrxn1α gRNA2 sequence is indicated by a solid black line, the protospacer adjacent motif (PAM) is indicated by a dashed red line, and the Cas9 cut site is indicated by a dashed black line.

(D) Quantification of targeting efficiencies for gRNAs against the three Nrxn1 alternative promoters and AS4 splice site in N2A-Cas9 by InDels.

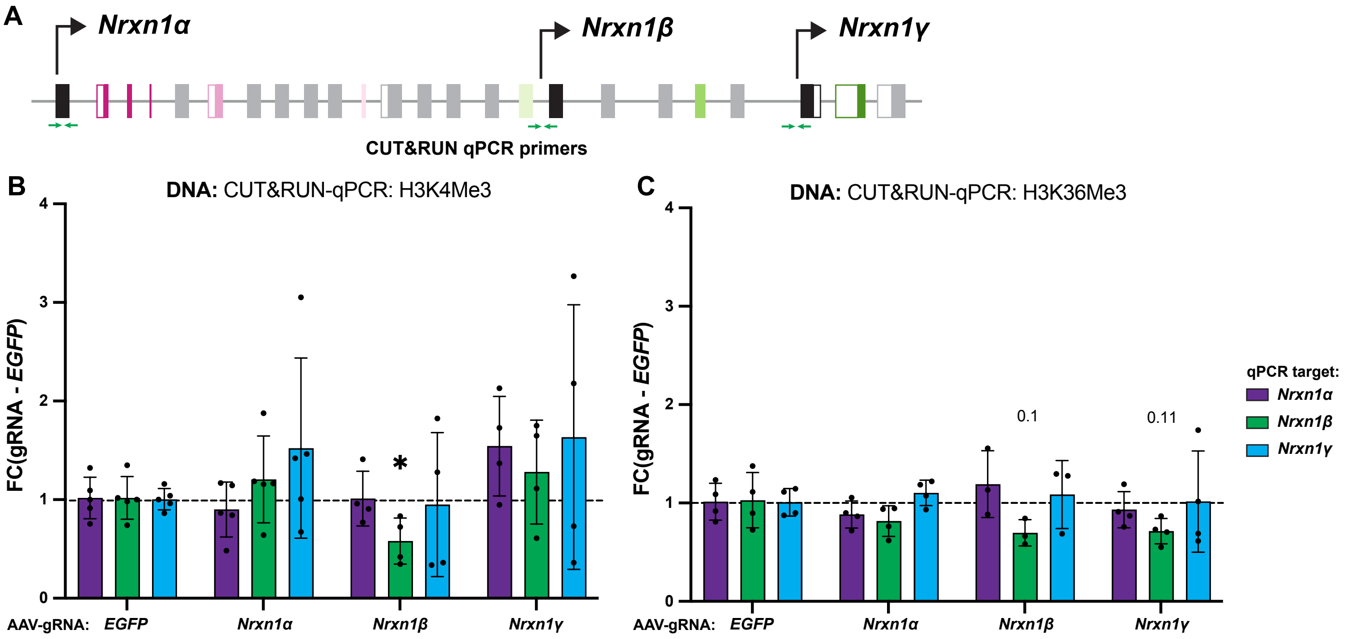


**Supplemental Figure 2. Epigenetics of transcriptional interference at Nrxn1 alternative promoters**
(A) Schematic of primers used to examine enrichment of histone modifications at individual alternative promoters by CUT&RUN-qPCR.
(B-C) CUT&RUN-qPCR against (B) H3K4Me3 and (C) H3K36Me3 for histone modifications around Nrxn1 alternative promoters in primary neurons generated from CRISPRi-KI mice after treatment with AAV-gRNAs. Enrichments of histone modifications around Nrxn1 alternative promoters were normalized to histone modifications around the Gapdh promoter. FC for each gRNA was calculated relative to a non-targeting EGFP gRNA. N=3-5 wells per group (* $p < 0.05$). p values are for gRNAs versus EGFP gRNA by unpaired student's t test.
